# DockQ v2: Improved automatic quality measure for protein multimers, nucleic acids, and small molecules

**DOI:** 10.1101/2024.05.28.596225

**Authors:** Claudio Mirabello, Björn Wallner

**Affiliations:** Division of Bioinformatics, Department of Physics, Chemistry and Biology, Linköping University, 581 83 Linköping, Sweden; National Bioinformatics Infrastructure Sweden, Science for Life Laboratory

## Abstract

It is important to assess the quality of modeled biomolecules to benchmark and assess the performance of different prediction methods. DockQ has emerged as the standard tool for assessing the quality of protein interfaces in model structures against given references. However, as predictions of large multimers with multiple chains become more common, there is a need to update DockQ with more functionality for robustness and speed. Moreover, as the field progresses and more methods are released to predict interactions between proteins and other types of molecules, such as nucleic acids and small molecules, it becomes necessary to have a tool that can assess all types of interactions. Here, we present a complete reimplementation of DockQ in pure Python. The updated version of DockQ is more portable, faster and introduces novel functionalities, such as automatic DockQ calculations for multiple interfaces and automatic chain mapping with multi-threading. These enhancements are designed to facilitate comparative analyses of protein complexes, particularly large multi-chain complexes. Furthermore, DockQ is now also able to score interfaces between proteins, nucleic acids, and small molecules.

**Code:** https://wallnerlab.org/DockQ

## Introduction

Protein interactions are crucial for most biological functions, including metabolism, gene expression, cell signaling, and immune response. Structural characterization of protein interactions at the molecular level is important to understanding basic biology, as well as disease mechanisms. Thus, elucidating protein structures has been the subject of immense research in previous decades. Experimental methods such as X-ray crystallography, NMR, and cryo-EM have determined the three-dimensional structure of over 200,000 proteins. However, experimental determination of protein structure is difficult, time-consuming, and sometimes even impossible given experimental constraints. Also, 200,000 structures are a small number compared to the over 250,000,000 proteins known to date. Thus, there has been a massive drive to develop computational methods to close this gap by predicting the structural properties of proteins.

To be able to compare the performance of computational methods it is important to have robust measures that can assess the quality of predicted models. Many such methods have been developed over the years, mostly for assessing monomer structures: MaxSub (1), GDT_TS (2), TM-score (3), and LDDT (4). Lately, methods have been developed also for multimers: DockQ (5), oligoLDDT (6), or US-align (7). In contrast to oligoLDDT and US-align, which measure the overall structural similarity of multimers, DockQ assesses the quality of the interactions between chains in multimers. This is an advantage since the quality of these interactions might be obscured in the overall structural similarity.

Lately, next-generation prediction tools, such as RoseTTAFold-All Atom (8) and AlphaFold 3 (9), have been released that can predict interactions between proteins, nucleic acids, and small molecules alike. In order to benchmark these new methods, it is important for a tool like DockQ to be able to evaluate interactions between all these types of molecules rather than forcing users to rely on multiple specialized tools.

We now introduce a new version of DockQ (v2). DockQ v2 is faster than the previous version, easier to install, and more portable. In this version, we include new functionalities: DockQ v2 can score multiple interfaces simultaneously in complexes with more than two chains and automatically detects the model-to-native chain mapping that maximizes the global quality of the complex. We also include the possibility of scoring interactions between proteins, nucleic acids, and small molecules. To the best of our knowledge, DockQ is the only tool that can seamlessly handle all these types of molecules.

In addition, we have also added interface clash detection, multithreading, transparent handling of mmCIF and gzipped files, the possibility to install it with *pip*, and the possibility to import it as Python module.

## Method description

### DockQ - assessing the quality of multimers

DockQ assesses the quality of a multimeric model by comparing it to a reference structure (typically an experimentally determined structure). The comparison is performed by assessing the similarity of the interfaces between protein chains. For a given protein-protein or protein-nucleic acid interface, the DockQ score is the average of the fraction of native contacts (*f*_*nat*_), scaled interface RMSD (iRMSD), and scaled ligand RMSD (LRMSD):

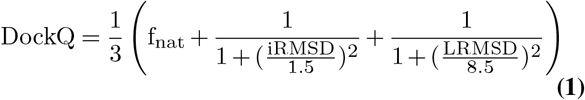

Where 1.5 and 8.5 are scaling parameters that monitors how fast iRMSD and LRMSD, respectively, should go to zero. These parameters were determined by maximizing the ability of DockQ to classify protein interfaces according to the CAPRI metrics (“Incorrect”, “Acceptable”, “Medium” and “High quality”) (10). The advantage of the DockQ score is that it is a continuous score in the [0, 1] range, from incorrect to high quality, which facilitates method comparisons since distributions, instead of classifications, of model quality scores can be compared. This also makes it better suited for machine learning (11).

To calculate the DockQ score using Eq. 1, the f_nat_, iRMSD, and LRMSD need to be calculated, below we describe how these measures are defined.

### Fraction of correctly predicted reference contacts

The f_nat_ measure is the fraction of correctly predicted reference or native contacts in the interface. An interface is a set of interacting amino acids or nucleotides sitting in separate chains with at least one pair of heavy atoms within 5Å of each other (4Å in the case of peptides). A contact is correct if a pair of interfacial amino acids in the model are also interacting in the reference. The total number of correct contacts is then divided by the total number of contacts in the model interfaces. A problem with f_nat_ is that it does not consider the number of false predictions. Thus, a model with a large number of false positives might still receive a fairly high f_nat_. Also, models with clashes will obtain higher f_nat_, taking it to the extreme, a model with all (*x, y, z*) coordinates at 0 will have f_nat_=1.0. This is, in general, only a problem for models with many clashes. Therefore, the number of clashes is automatically detected and reported to the user in DockQ v2.

### Interface RMSD

The interface RMSD, iRMSD, is the backbone RMSD of the interface residues. Interface residues are defined as pairs of residues between protein and/or nucleic acid chains where any two heavy atoms are within 10Å of each other (8Å between pairs of *Cβ* atoms in case of peptides).

### Ligand RMSD

The ligand RMSD, LRMSD, is the backbone RMSD of the ligand when superimposing on the receptor. The receptor is defined as the larger of the two chains by number of amino acids or nucleotides. If two chains are equal in size, the first will be considered the receptor. In the case of small molecule ligands, LRMSD is calculated on the position of all heavy atoms in the ligand when superimposing on the receptor interface (*pocket-aligned LRMSD*).

### Workflow and new functionalities

DockQ used to be protein-only. We have now added the functionality to also score interactions involving nucleic acids, and small molecule ligands. In the case of small molecule ligands, only the pocket-aligned ligand RMSD (LRMSD) is reported, as commonly done in other methods (12, 13) and the corresponding DockQ score is calculated only based on the LRSMD component in Eq. 1 (f_nat_ = *iRMSD* = 0). Since small molecules are not ordered like proteins and nucleic acids, we use NetworkX’s (14) graph matching functionality and report the solution with lowest LRMSD when multiple matches are possible (e.g. symmetric molecules). Graph edges are calculated according to the atoms’ covalent radii (15).

DockQ reads the coordinates of a model and reference structure in PDB or mmCIF format, either uncompressed or compressed with *gzip*, to calculate the DockQ score (Eq. 1) for each interface in the reference structure. It returns the DockQ score for each interface as well as an overall *GlobalDockQ*, defined as the average of DockQ scores over all interfaces. Equivalent residues in the model and reference are mapped using sequence alignment (default) or by following the residue numbering.

For multimeric complexes with subunit stoichiometry larger than one, there is a need to find the optimal chain mapping, i.e. the chain mapping that maximizes the overall GlobalDockQ. This is also needed when multiple small molecules of the same type are being evaluated at once. The number of possible chain mappings grows with the factorial of the number of equivalent molecules, i.e., a homodimer of stoichiometry A2 has 2! = 2 possible mappings, while a homooctamer of stoichiometry A8 has 8! = 40, 320. The default behavior is that DockQ will identify equivalent subunits by sequence alignment (by molecule name in the case of small molecule ligands) and exhaustively permutate the chain mappings to find the one with the highest GlobalDockQ. It is also possible to provide a given chain mapping with the *−− mapping* option or to limit the search to a subset of native chains. The search is parallelized over multiple threads (default=8) using LRU caching and with computationally intensive functions, such as distance calculations, implemented in Cython for speed.

DockQ v2 is released as a pip-installable Python package. This allows you to import and use DockQ as a module in other software.

## Results

To verify that DockQ v2 is able to reproduce the DockQ scores generated with the previous version, we compared DockQ scores calculated for 17,409 models from 37 CASP15 targets. The agreement is almost perfect (R=1.000) between the new and the previous version; see Figure 1a. Thus, the scores from both versions can be used interchangeably.

**Fig. 1.**
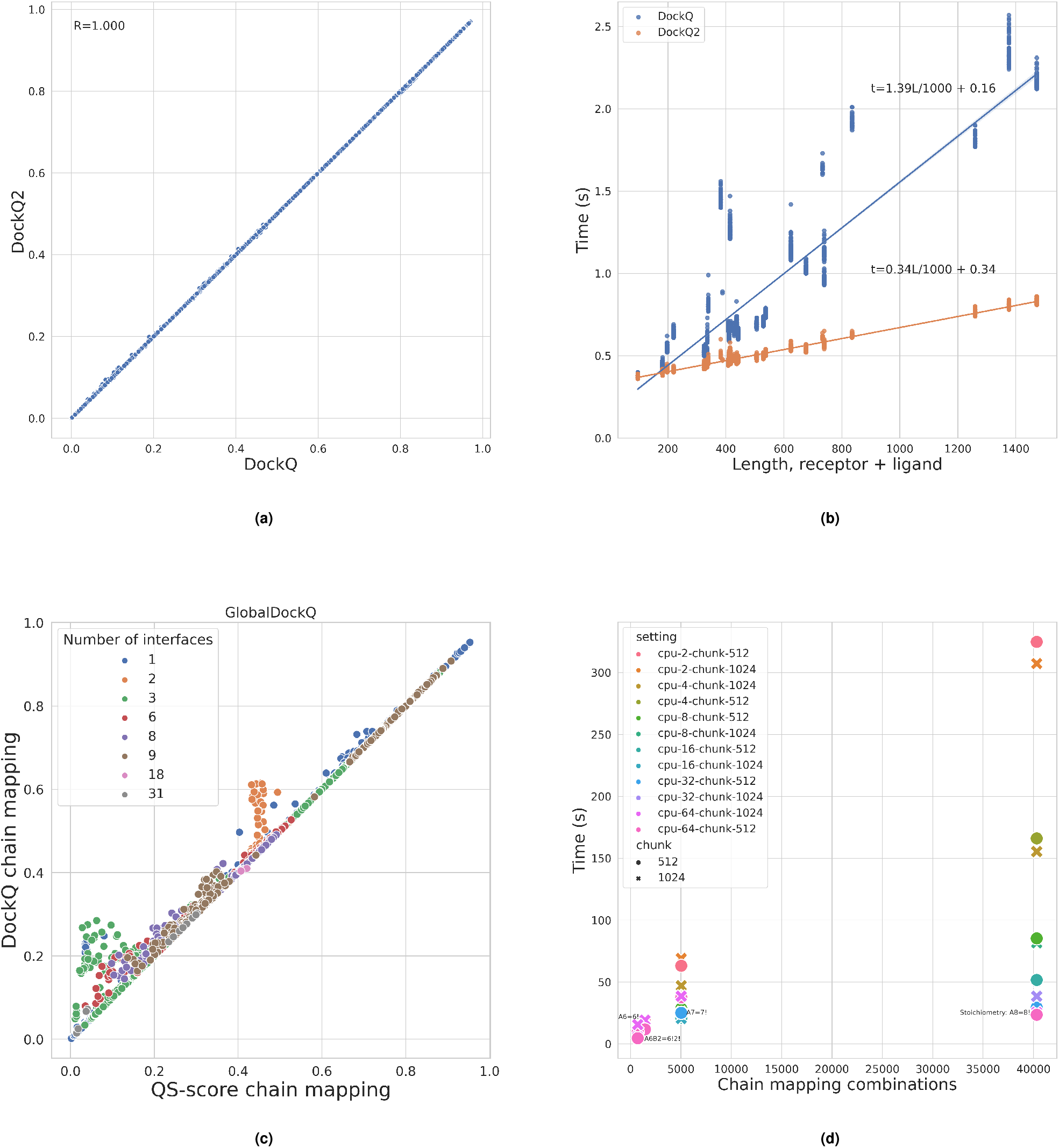
(a) We evaluated DockQ v2 against the old DockQ (v1.0) on a large benchmark of 17,049 models from 37 CASP15 targets. The DockQ scores are equivalent between the two versions. (b) The new version of DockQ is faster than the previous due to optimised implementation of computationally expensive functions in Cython, especially for larger complexes. (c) Comparison of methods to perform automatic mappings between model and native chains: the automatic mapping done in DockQ yields the same or better GlobalDockQ than when using mappings optimised by QS-score. (d) Time to exhaustively optimise the mapping between sets of chains for complexes with different stoichiometry. The time complexity is linear with the number of chain mapping combinations, and can be proportionally cut down by increasing the number of parallel threads in DockQ.

The updated version is generally faster, especially for larger targets, as shown in Figure 1b. The linear portion of the time complexity is decreased approximately four times (4.08) compared to the previous version. This leads to significantly reduced runtimes in practice.

An important new feature is automatic chain mapping, which simplifies the problem of finding the optimal chain mapping between symmetric chains. Previously, the optimal chain mapping has either been ignored, assuming a one-to-one correspondence between the chains in model and reference, or had to be determined with some other software like the chain mapping routine in QS-score (16). The latter was used when calculating DockQ in the evaluation of multimer predictions in CASP15 (17). We compared the DockQ scores obtained using the new automatic chain mapping to the DockQ scores obtained by using the mapping from the chain mapping routine in QS-score. In all cases, the automatic chain mapping in DockQ produces an identical or higher GlobalDockQ compared to using the mapping provided by QS-score, Figure 1c. This is expected since the mapping using QS-score is optimized using QS-score, while the automatic chain mapping in DockQ is optimized on GlobalDockQ. To make the automatic chain mapping faster, it is parallelized on multiple threads, making it possible to exhaustively map a homomer with stoichiometry A8 (8! = 40, 320 chain combinations) in less than a minute on 8 threads or one with stoichiometry A7 (7! = 5040 combinations) in less than 30 seconds, see Figure 1d.

Taken together, we believe that this new, improved version of DockQ will become an essential tool for evaluating models of biomolecular complexes, regardless of the type of molecule they may include.

## ACKNOWLEDGEMENTS

This work was supported by the Wallenberg AI, Autonomous System and Software Program (WASP) from Knut and Alice Wallenberg Foundation (KAW), Swedish Research Council grant, 2020-03352, The Swedish e-Science Research Center, and Carl Tryggers stiftelse för Vetenskaplig Forskning, 20:453. CM is financially supported by the Knut and Alice Wallenberg (KAW) Foundation and the “BeyondFold” Technological Development Project at SciLifeLab. The computations were performed on resources provided by KAW and NSC (Berzelius).

